# Resolving Conformational Heterogeneity in Intrinsically Disordered Proteins via Experimentally Guided Multi-Replica Simulations

**DOI:** 10.64898/2026.01.29.702558

**Authors:** Wangfei Yang, Sichun Yang, Wenwei Zheng

## Abstract

The conformational landscapes of intrinsically disordered proteins (IDPs) remain obscured by the ensemble averaging inherent to experimental observables. Here, we present Multi-replica Averaged Restraint Simulation (MARS), a data-driven modeling framework that reconstructs these landscapes by enforcing ensemble-averaged restraints across multiple replicas without imposing prior structural knowledge. Using the N-terminal domain of estrogen receptor alpha (ERα-NTD) as a model system, MARS simultaneously integrates small-angle X-ray scattering (SAXS) and six paramagnetic relaxation enhancement (PRE) profiles, comprising over 600 pairwise restraints, to generate ensembles that quantitatively fit all input structural restraints. The resulting ensemble is consistent with independent backbone relaxation measurements and reveals two major conformational states: a dominant extended state and a low-populated yet functionally relevant compact state whose structural features align with prior mutagenesis studies. Systematic benchmarking demonstrates that SAXS and PRE provide orthogonal global and local constraints, that each PRE profile contributes non-redundant structural information, and that multi-replica sampling is essential for preserving conformational heterogeneity. MARS offers a scalable framework for integrating orthogonal biophysical measurements to resolve both dominant and rare functional states in IDPs.

## Introduction

Intrinsically disordered proteins (IDPs) constitute a substantial proportion of the proteome and play essential roles in diverse biological processes^1^. However, their structural flexibility makes them more challenging to study than folded proteins. Unlike folded proteins, whose relatively stable three-dimensional structures can be directly determined^2,3^, IDPs lack a fixed conformation and instead interconvert dynamically among a vast ensemble of structures^4^. Experimental measurements therefore report ensemble averages over all accessible conformations, obscuring the underlying conformational heterogeneity, particularly rare conformations that may be functionally important. Resolving these hidden states from ensemble-averaged experimental observables remains a central challenge in IDP structural characterization.

In this context, computational modeling is indispensable for constructing conformational ensembles of IDPs and interpreting experimental observations. Over the past several decades, approaches such as Monte Carlo (MC) sampling^5,6^ and molecular dynamics (MD) simulations^7^ have been widely used to generate conformational ensembles of IDPs (at both all-atom^8–10^ and coarse-grained^11–13^ resolutions). More recently, advances in machine learning have led to the application of data-driven techniques to IDP ensemble construction. For example, inspired by the success of AlphaFold for folded proteins^14^, generative models have been developed to directly produce IDP conformational ensembles^15–18^, and machine-learning force fields have also been integrated with traditional simulation frameworks^19–23^. However, achieving consistent agreement with diverse experimental measurements remains challenging, highlighting the importance of understanding the complementary structural information provided by different experimental techniques.

Although the conformational ensembles of IDPs cannot be directly determined, experiments provide access to structural features spanning different length scales and resolutions. For example, the radius of gyration (*R*_*g*_), which characterizes the overall dimensions of an IDP, can be measured by Small-Angle X-ray Scattering (SAXS)^24^; distances between specific residue pairs can be obtained from Förster Resonance Energy Transfer (FRET)^25^; and local properties such as secondary structure propensities can be inferred from Nuclear Magnetic Resonance (NMR) spectroscopy^26^. In addition, more detailed multi-distance information can be extracted from paramagnetic relaxation enhancement (PRE) measurements^27^. These experimental observables impose increasingly stringent restraints on IDP conformational ensembles^28^.

Among recent computational approaches that integrate physics-based models with experimental data to generate structural ensembles of IDPs, ensemble reweighting^29–34^ and Bayesian ensemble refinement^35–37^ are commonly used strategies. These posterior methods adjust a pre-generated conformational pool to match experimental observables and have seen broad success, including in studies combining SAXS, FRET and NMR data.^38–45^ Their effectiveness nonetheless depends on the completeness of the initial pool, as increasingly informative experimental datasets expose limitations intrinsic to incomplete sampling and inaccuracies in the underlying physics-based models. An alternative class of approaches incorporates experimental data directly during conformational sampling through biasing terms added to the potential energy. While such restraint-based simulations are routinely used for folded proteins^46–48^, their extension to IDPs is complicated by the breadth and heterogeneity of IDP conformational ensembles. Multi-replica schemes have therefore been developed to enforce agreement with ensemble-averaged experimental properties^49– 52^. However, the practical requirements and performance limits of these approaches, particularly for integrating diverse experimental measurements in IDPs, remain poorly understood.

In this study, we introduce Multi-replica Averaged Restraint Simulation (MARS), an experimental data–restrained simulation framework for IDPs, which applies experimental restraints to ensemble-averaged observables across multiple parallel replicas rather than to individual trajectory. Using the N-terminal domain of estrogen receptor α (ERα-NTD) as a model system^53^, we demonstrate that MARS preserves conformational heterogeneity while steering sampling toward simultaneous agreement with diverse experimental measurements, including SAXS data and PRE profiles from six spin-labeling positions comprising more than 600 pairwise restraints^53^. The resulting conformational landscape is in close agreement with experimental observations and resolves two major conformational states, including a low-populated yet functionally relevant compact state, consistent with prior mutagenesis studies^53^. We further systematically benchmark the influence of replica number and restraint composition on ensemble quality, providing practical guidance for applying ensemble-averaged restrained simulations to IDPs.

## Results and Discussion

### Implementation and Validation of MARS using ERα-NTD

MARS integrates experimental restraints by enforcing agreement between ensemble-averaged observables and experimental data across multiple replicas in MD simulations. The workflow begins with the initiation of multiple parallel simulations. At each simulation step, experimental observables are evaluated across all replicas, and harmonic potentials are applied according to deviations between ensemble-averaged values and the corresponding experimental measurements (**Figure 1**). The resulting conformational ensemble, composed of multiple simulation trajectories, is then analyzed to extract relevant molecular determinants.

**Fig. 1.**
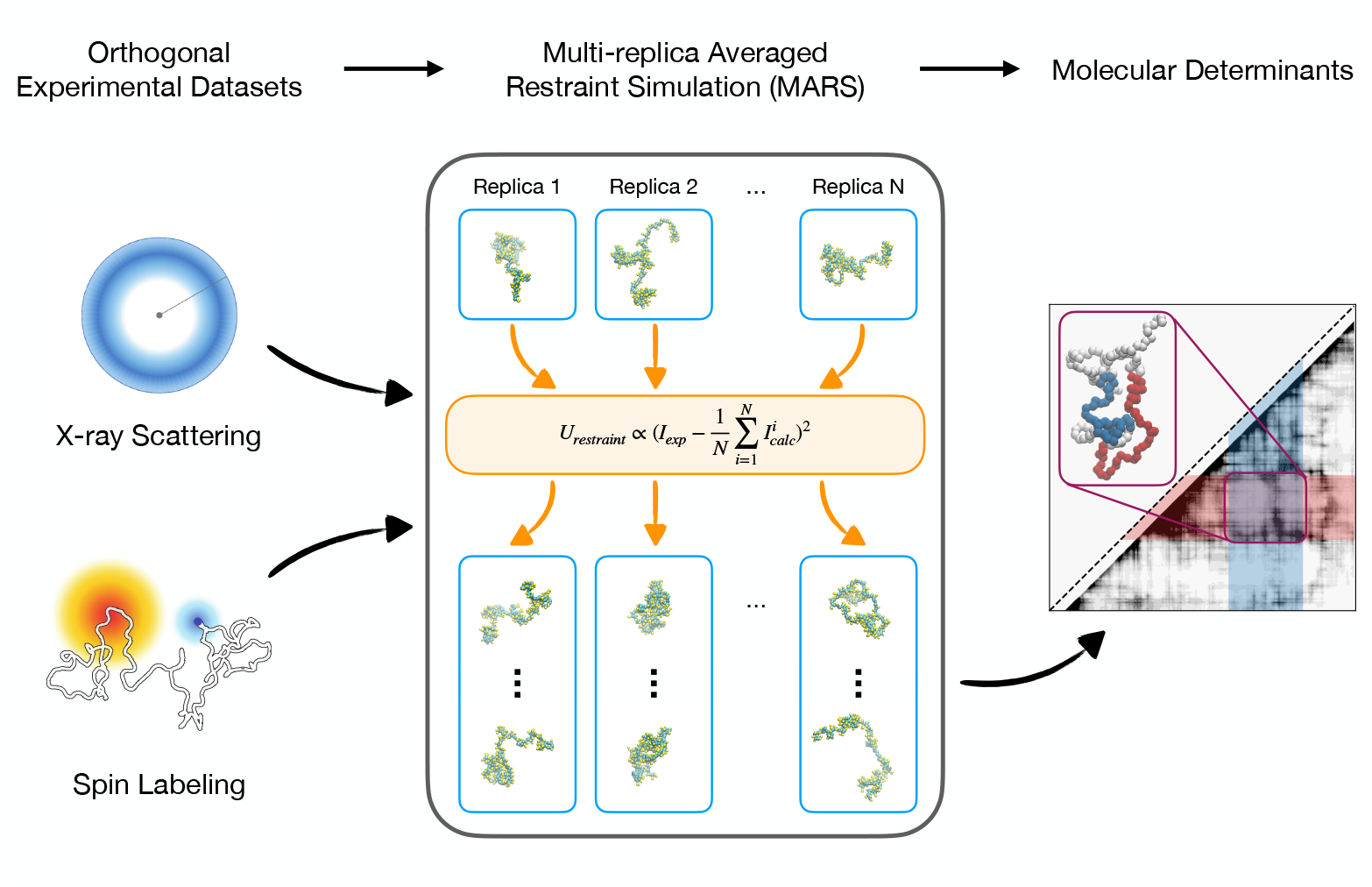
The MARS workflow for constructing IDP conformational ensembles. Left: Orthogonal experimental datasets serve as restraints for constructing conformational ensembles, integrating both global properties, such as overall protein dimensions from SAXS, and local features, such as site-specific distances between spin labels (orange) and backbone atoms (blue) from PRE. Middle: MARS executes multiple parallel replica simulations, with experimental restraints simultaneously applied to ensemble-averaged observables across all replicas. Right: Molecular determinants, such as residue-based contact maps, can be extracted from the resulting conformational ensembles.

To evaluate how effectively this strategy integrates orthogonal measurements for IDPs, we applied our framework to ERα-NTD, the intrinsically disordered regulatory region of ERα that contributes to ligand-independent transcriptional activation via a hydrophobic-driven mechanism involving two aromatic-rich sequence clusters (residues 43-81 and residues 108-152).^53^ This system is well suited for benchmarking because of extensive available experimental data, including SAXS profiles that report on global dimensions and PRE profiles from six evenly-distributed labeling positions along the sequence that provide sensitive measures of short-to intermediate-range residue-residue interactions.

We implemented MARS using the coarse-grained MARTINI2 model^54^, where each residue is represented by one to five beads and four water molecules are mapped as one single bead, in combination with PLUMED^55^ to apply experimental restraints. We ran 32 parallel simulations; at each simulation step, SAXS intensities and PRE intensity ratios were computed for every replica and averaged across the ensemble. Restraint potentials were then applied independently to the SAXS and PRE observables. Because SAXS and PRE arise from distinct physical principles and exhibit different dependencies on pairwise distances, ensemble averaging was performed using observable-specific schemes to ensure proper application of the biasing potentials (see Eqs. 1 and 2). Although demonstrated here with a coarse-grained model, the MARS workflow is generally applicable to any physics-based simulation model, including all-atom explicit-solvent simulations. Further details of potential forms, strength tuning, and simulation setup are provided in Methods and **Figures S1** and **S2**.

Simulation convergence was assessed by analyzing the time autocorrelation of the end-to-end distance. The corresponding relaxation time was approximately 40 ns (**Figure S2C**), more than an order of magnitude shorter than the total simulation length of 500 ns and comparable to experimentally reported chain relaxation times for IDPs^56^. In addition, the *R*_*g*_ distributions from independent replicas exhibited substantial overlap and broad sampling, indicating consistent convergence of the simulations across multiple replicas (**Figure S2D**). Accordingly, the first 100 ns of each simulation were discarded as equilibration, and the remaining 400 ns were used for subsequent analysis.

Using MARS we obtained a conformational ensemble for ERα-NTD in excellent agreement with experimental measurements (**Figure 2**). Following standard practice, we quantified deviations between simulation and experiment using the *χ*^2^ statistic for SAXS and the *Q* factor for PRE (see Methods). For SAXS, we obtained a *χ*^2^ value of 1.2 (**Figure 2A**); as direct validation, the *R*_*g*_ determined from SAXS data was 33.9 Å, in close agreement with the ensemble-averaged value of 33.2 Å from our simulations. For the six PRE profiles, *Q* factors ranged from 0.11 to 0.17 (**Figure 2B**), indicating excellent agreement. Together, these metrics demonstrate that the MARS-derived ensemble quantitatively reproduces both global and local experimental observables.

**Fig. 2.**
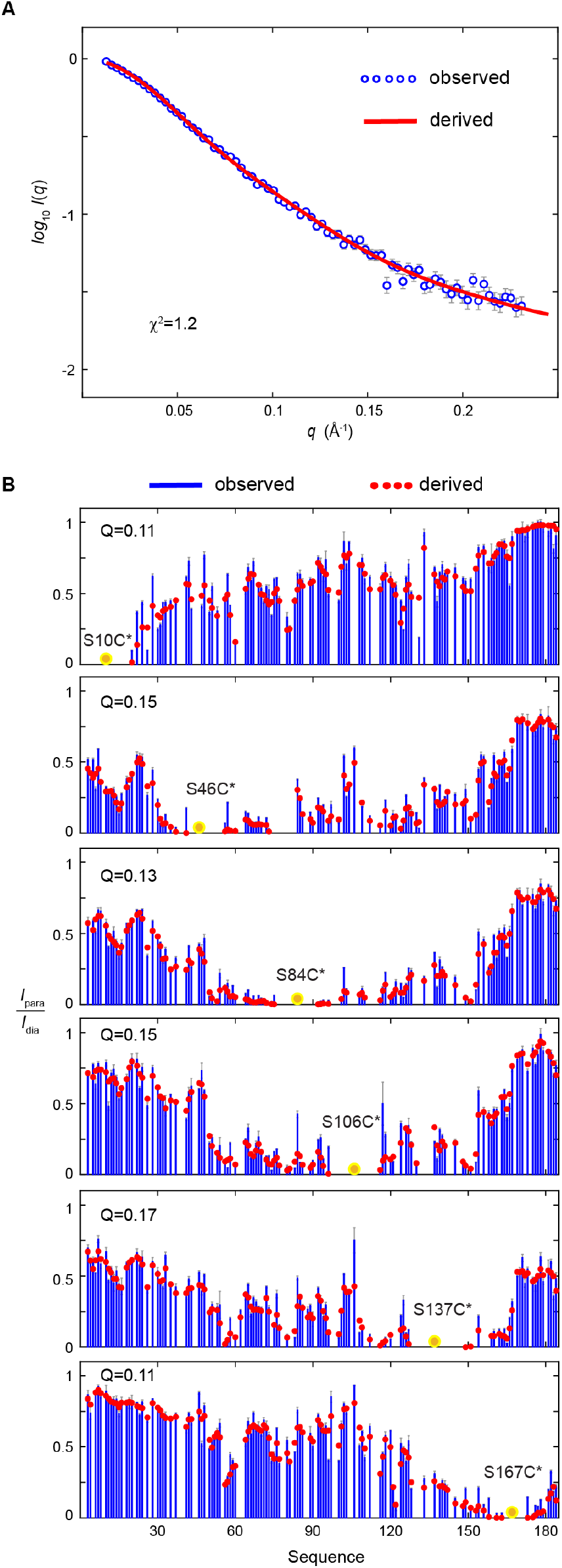
Simultaneous matching of SAXS and PRE experimental data by MARS simulations. **(A)** Comparison of experimentally-observed (blue) and simulation-derived (red) SAXS intensity profiles (*χ*^*2*^ = 1.2). *I(q)*, normalized scattering intensity; *q*, scattering vector amplitude. **(B)** Comparison of observed (blue) and derived (red) PRE profiles expressed as peak intensity ratios (*I*_para_/*I*_dia_) for six MTSL-labeled single-cysteine variants (S10C, S46C, S84C, S106C, S137C, and S167C). Yellow circles indicate spin-label positions. *Q* factors for each profile are shown in the legend. Experimental SAXS and PRE data were obtained from the previous literature.^53^

To assess whether MARS overfits the SAXS and PRE datasets, we evaluated several other experimental observables that were not included in the restraints. Chemical shifts calculated from the resulting ensemble using SPARTA+^57^ showed close agreement with experimental measurements (**Figure S3**), demonstrating that the ensemble captures local backbone environments without being explicitly restrained. We also examined NMR relaxation parameters, which report on backbone dynamics across multiple timescales. Because hydrogen atoms were not explicitly represented in the simulations, the absolute timescales of the computed relaxation parameters are not expected to match those measured experimentally. Nevertheless, residues predicted by the ensemble to experience greater local flexibility or transient ordering showed patterns consistent with published relaxation measurements (**Figure S4**).^53^ Together, these cross validations indicate that MARS produces physically realistic ensembles rather than reproducing the input data through overfitting.

### The Resulting Conformational Landscape Reveals Two Major States

Beyond ensemble-averaged properties, we examined the underlying conformational landscape to identify the dominant slow motions governing its dynamics. Due to the inherent flexibility of IDPs, extracting meaningful information from high-dimensional trajectory data using conventional methods such as root-mean-square-deviation (RMSD) clustering is challenging. To address this, we employed time-lagged independent component analysis (TICA), a dimensionality reduction method for time-series data that incorporates temporal correlations.^58^ Briefly, TICA identifies dominant slow motions by solving the eigenvalue problem of the time-lagged covariance matrix under the assumption of Markovian dynamics, yielding collective variables termed TICA coordinates (see Methods). Using TICA, we identified a small number of dominant slow motions, represented by the leading TICA coordinates, with associated timescales shown in **Figures S5A** and **S5B**.

Projections of converged MARS-generated ensemble, composed of 32 replicas of 400 ns each, onto the leading TICA coordinates (TICA1 and TICA2) revealed two distinct conformational states (**Figure 3A**). The boundary of each state was defined by applying a log-probability threshold of 1.0 *k*_B_T. The extended state accounted for 46.6% of the ensemble, whereas the compact state represented only 8.4%. Representative structures, selected as those with the lowest RMSD relative to all other structures within the same state, are displayed alongside the contact maps. When SAXS and PRE observables were computed separately for the extended and compact states (**Figure S6**), neither state alone reproduced the experimentally averaged measurements, indicating that the experimental data reflect a conformationally heterogeneous ensemble rather than a single dominant structure. Experimentally, low-populated states are difficult to characterize because they are largely obscured by the dominant population. However, such states can play crucial roles in biological function. For example, extensive mutagenesis studies have demonstrated that the compact state of ERα-NTD is essential for its phosphorylation-dependent transcriptional activity.^53^

**Fig. 3.**
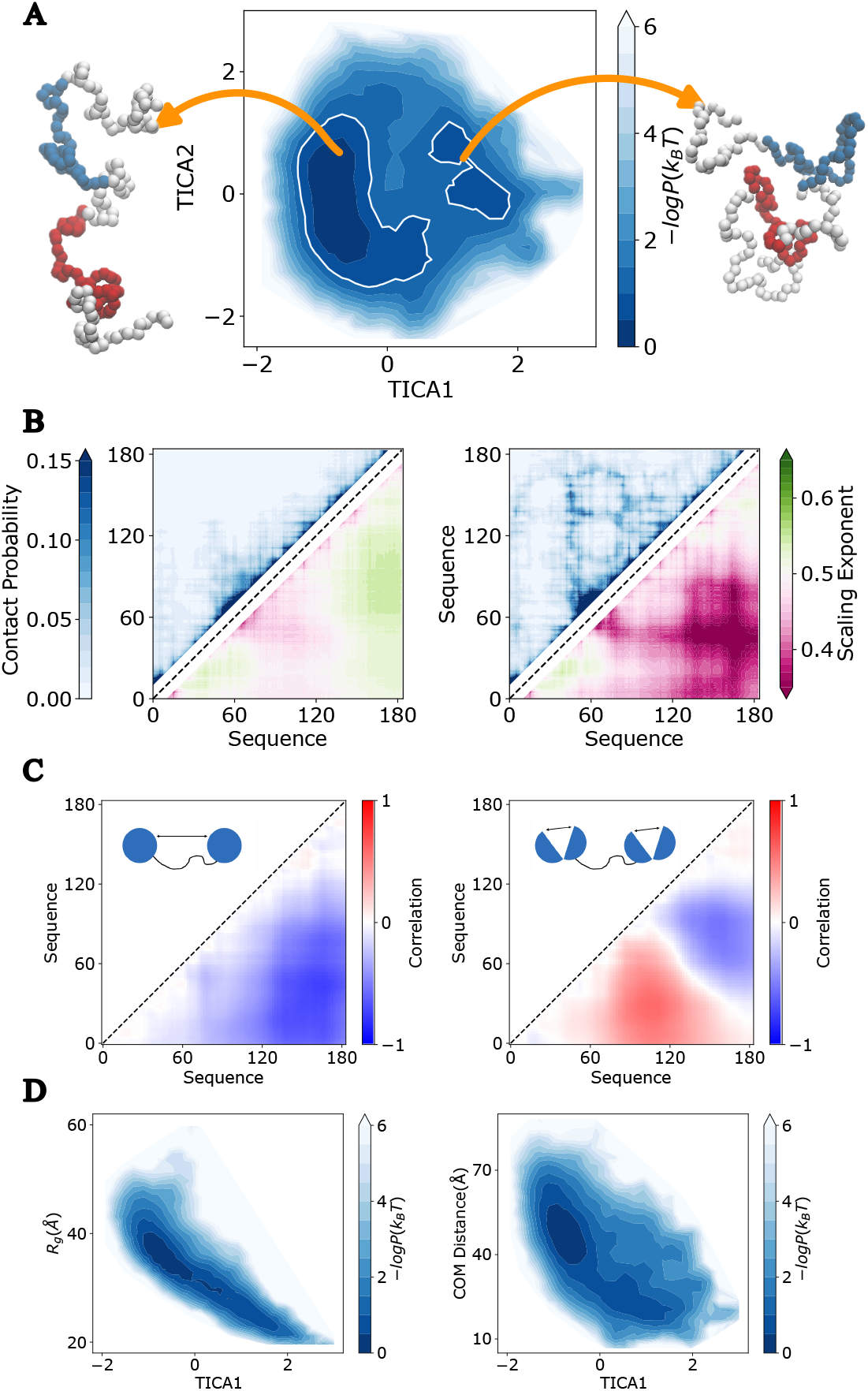
Conformational landscape of ERα-NTD. **(A)** Projection of conformational landscape onto the leading TICA coordinates (TICA1 and TICA2), corresponding to the slowest motions, with two conformational states delineated by a log-probability threshold of 1.0 *k*_B_T (white curve). Representative conformations from each state (compact and expanded) are shown on each side. **(B)** Contact map (upper diagonal) and scaling exponent map (lower diagonal) for the two conformational states. **(C)** Correlations between pairwise distances and the TICA1 (left) and the TICA2 (right) coordinates, with illustrations of the corresponding motion modes shown in the upper diagonal panel. **(D)** Conformational landscape projected along TICA1 versus *R*_*g*_ and COM distance between the two aromatic-rich sequence clusters (residues 43-81 and residues 108-152).

To gain structural insight into the differences between the identified states, we analyzed residue– residue contact maps, which provide a compact representation of intramolecular interactions. Contact maps computed using a 12 Å distance cutoff between the closest beads (**Figure 3B**, upper diagonal) reveal that the extended state is dominated by local interactions between sequentially adjacent residues, whereas the compact state exhibits pronounced long-range interactions. This result is consistent with previous experimental observations demonstrating that ERα-NTD can adopt a more closed conformation through interactions between two aromatic-rich clusters in the absence of serine 118 phosphorylation.^53^ Indeed, in the compact state, interactions between these two aromatic-rich clusters are evident.

While contact maps primarily reports on short-range interactions, they do not directly capture how the chain organizes over large sequence separations. To characterize this long-range chain behavior, we examined polymer scaling exponents ν, which describe how intrachain distances grow with sequence separation. Accordingly, we computed residue-pair scaling exponents *v*_*i,j*_ = In(*r*_*i,j*_/*b*) /ln (|*i* − *j*|), in which *r*_*i,j*_ is the distance between the *i*-th and *j*-th amino acids and *b*=5.5Å.^59,60^ In polymer physics, the expected value of *v* is 0.5 for an ideal Gaussian chain and approximately 0.588 for a self-avoiding chain in the excluded-volume limit, providing a reference for interpreting local chain compaction or expansion.^61^ In the scaling exponent map (**Figure 3B**, lower diagonal), interactions between the aromatic-rich clusters become even more pronounced, with scaling exponents between the two clusters dropping below 0.4. Such reduced *v* values indicate strong effective compaction relative to a random coil, consistent with persistent contacts that locally collapse the chain across long sequence separations.

Because TICA coordinates themselves are abstract collective variables, their physical interpretation requires connecting them back to the underlying molecular degrees of freedom. In our analysis, TICA was performed with pairwise distances between residues sampled at alternating sequence positions. To interpret the leading TICA modes, we examined how these pairwise distances correlate with the first two TICA coordinates (**Figure 3C**). The resulting matrices quantify the correlation between each individual pairwise distance and a given TICA coordinate. Larger absolute correlation values indicate that variations in a specific pairwise distance contribute to the associated slow TICA collective motion.

Using these weights, we found that TICA1, the first TICA coordinate corresponding to the slowest motion, is dominated by inter-cluster distances between the two aromatic-rich clusters, which primarily determines the overall molecular size. Consistent with this interpretation, the free-energy landscape projected onto TICA1 and either the *R*_*g*_ or the center-of-mass (COM) distance between the two clusters shows strong correlations between these metrics (**Figure 3D**), demonstrating that the two conformational states are largely distinguished by global chain dimensions. This behavior is consistent with the open–closed mechanism for ERα-NTD inferred from previous experimental studies. However, this correlation weakens at moderate to large *R*_*g*_ values, suggesting that *R*_*g*_ alone does not fully capture the internal conformational variability of the extended state and may not serve as a reliable reaction coordinate for IDPs.

In contrast, TICA2, the second TICA coordinate corresponding to the second slowest motion, is primarily influenced by intra-cluster distances, reflecting internal rearrangements within each aromatic-rich cluster (**Figure 3C**). The free-energy landscape projected onto TICA2 reveals substantial conformational variability within the intermediate region connecting the extended and compact states (**Figure 3A**), and these internal rearrangements occur on a slightly faster timescale than global compaction (**Figure S5B**). Notably, free-energy projections of TICA2 against either the *R*_*g*_ or the inter-cluster COM distance show no clear correlation (**Figure S5C** and **S5D**), indicating that TICA2 captures internal motions largely decoupled from global chain dimensions. As a result, TICA2 complements TICA1 by capturing heterogeneity along the transition between metastable states, suggesting the existence of alternative pathways in which global chain collapse is coupled to distinct sequences of intra-cluster reorganization rather than proceeding through a single well-defined route.

Together, these analyses yield a low-dimensional representation of the conformational landscape of the MARS-constructed ensemble, revealing two dominant conformational states of ERα-NTD separated along the slowest collective modes. Notably, the low-populated compact state is consistent with prior mutagenesis studies indicating functional relevance, yet would remain obscured in ensemble-averaged observables without the use of restraint simulations. Further analyses using contact and scaling exponent maps indicate that this compact state is stabilized by persistent inter- and intra-cluster interactions, whereas the dominant extended state lacks significant long-range interactions. Overall, the combination of MARS and TICA-based analysis enables direct resolution of the slow collective motions that distinguish these conformational states.

### Both Orthogonal Experiments and Multi-replica Restraints are Indispensable to Resolve IDP Conformational Heterogeneity

Beyond demonstrating the performance of MARS, we addressed broader questions in the IDP field concerning the experimental inputs and modeling strategies required to resolve conformational heterogeneity. Specifically, we evaluated: (i) the performance of traditional posterior fitting approach on the same task, (ii) the complementary roles of SAXS and PRE in constraining global and local conformational features, and (iii) the number of replicas needed to capture the ensemble-averaged nature of experimental observables. Together, these analyses provide a controlled assessment of the methodological requirements for reconstructing IDP conformational ensembles and clarify why orthogonal experimental datasets and multi-replica restraints are essential for resolving conformational heterogeneity.

To assess the performance of traditional posterior fitting, we first simulated ERα-NTD without experimental biases, using the same simulation length and number of replicas to generate the initial conformational pool. Conformation weights were then iteratively adjusted to minimize the Root Mean Square Error (RMSE) of calculated SAXS and PRE data from the experimental data, with a Shannon entropy term included to limit deviation from the original ensemble and avoid overfitting (see Methods and **Figure S7**). While ensemble reweighting successfully reproduced SAXS intensities, it failed to achieve close agreement with PRE (**Figure 4A** and **Figure S7**). Projection of the reweighted ensemble onto the TICA space of the MARS-derived ensemble reveals a fundamental limitation of posterior fitting: because reweighting only redistributes weights among pre-existing conformations, it cannot recover structures absent from the initial pool (**Figure 4B**). As a result, its performance depends critically on the ability of the underlying physics-based model to sample all experimentally relevant conformations in advance, a requirement that is challenging even by using all-atom force fields^63^, relative to the coarse-grained MARTINI ensemble used here, when local restraints such as PRE are involved.

**Fig. 4.**
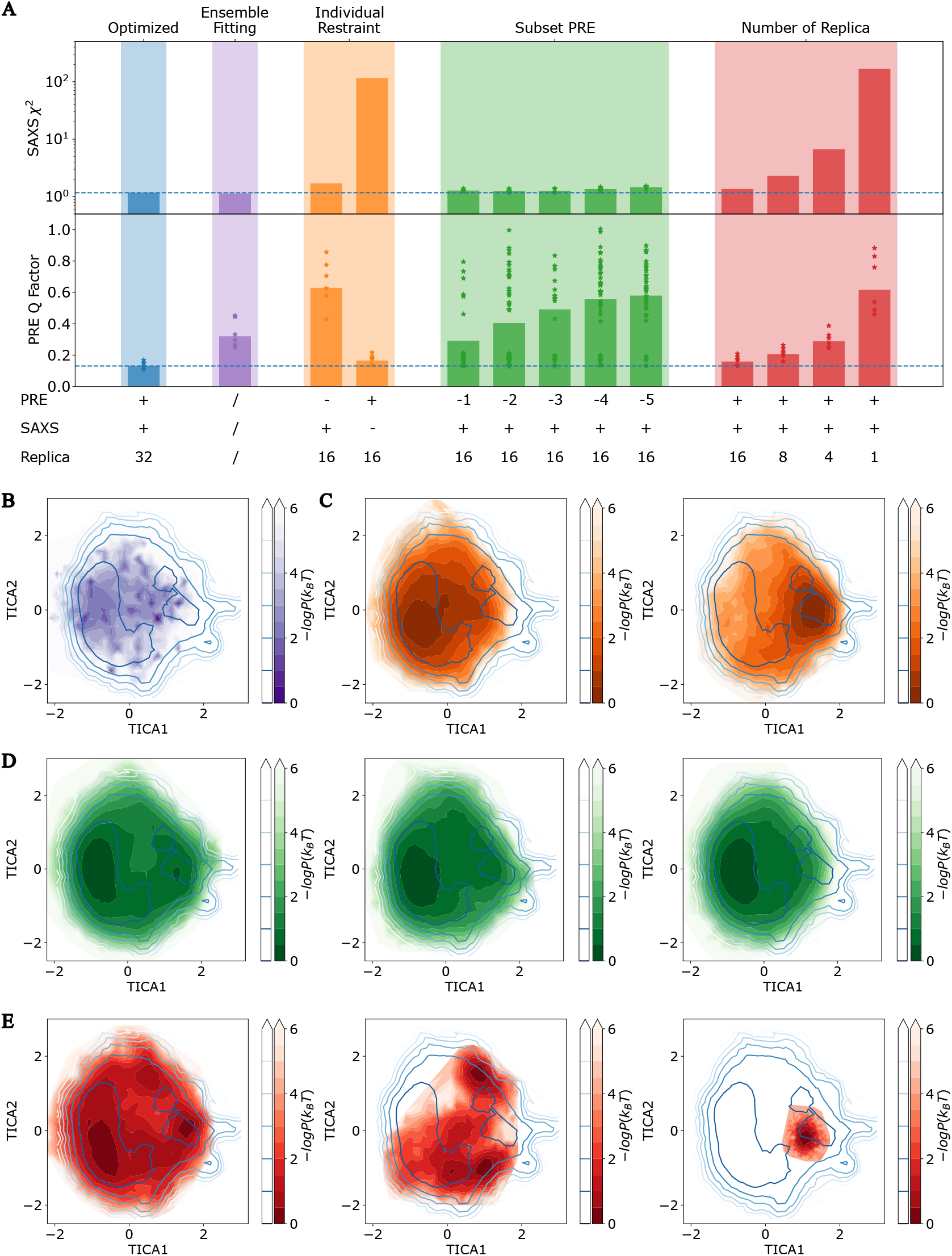
Dissecting the Contributions of Experimental Restraints and Replica Sampling to Ensemble Construction. **(A)** Agreement with experimental SAXS and PRE measurements for various ensemble construction methods, including: the optimized MARS pipeline with both SAXS and PRE restraints and 32 replicas (blue), simulations restrained by SAXS or PRE alone (orange), simulations restrained by SAXS with a reduced number of PRE profiles (green; labels indicate the number of PRE profiles used), simulations using fewer replicas under combined SAXS and PRE restraints (red; labels indicate the number of replicas), and traditional posterior ensemble reweighting method (purple). Stars indicate the Q factors of individual PRE profiles. **(B)** Conformational landscape projected onto the TICA space for the ensemble obtained by traditional posterior ensemble reweighting method. **(C)** Conformational landscape for simulations restrained by only SAXS (left) or PRE data (right). **(D)** Conformational landscape for simulations restrained by SAXS and five (left), three (middle), or one (right) PRE profiles. **(E)** Conformational landscape for simulations restrained by SAXS and PRE using 16 (left), 4 (middle), or 1 (right) replica.

To examine the interplay between the global and local information provided by SAXS and PRE, we performed individual restraint simulations in which either the SAXS or PRE restraints were removed from the original setup. The two datasets were found to be orthogonal and strongly complementary: removing either restraint led to a substantial loss of agreement with the corresponding experimental data (**Figure 4A** and **Figure S8**). This behavior can be explained by two factors. First, both global and local observables are inherently degenerate: conformations consistent with SAXS-constrained global dimensions can exhibit diverse local contact patterns, while conformations satisfying specific PRE distance restraints can span a broad range of overall sizes. Second, SAXS and PRE weight conformations differently across spatial scales: all conformations contribute equally to SAXS intensity, whereas PRE signals are dominated by a small subset of conformations in which specific residue pairs transiently approach. Consequently, even a minor population of such conformations can reproduce PRE measurements. To further illustrate this complementarity, we projected trajectories from the single-source simulations onto the same TICA space as the original simulation. The results confirm that SAXS and PRE restraints constrain the ensemble along complementary directions of TICA1 (**Figure 4C**), with SAXS favoring extended global dimensions and PRE favoring local proximity between specific residues.

Although the PRE dataset contains a large number of restraints, it is unclear whether the six PRE profiles provide redundant or complementary information. To address this, we performed a jackknife analysis in which one to five PRE profiles were selectively removed while all other aspects of the simulations, including SAXS restraints, were kept unchanged. For cases involving two, three, or four PRE profiles, where the number of possible combinations is large, a representative subset of combinations was evaluated, as summarized in **Table S1**. If the dataset were overdetermined, removing profiles would not substantially degrade agreement across all six. However, removing even a single PRE profile increased the overall *Q* factor, with the increase almost entirely attributable to the removed profile (**Figure 4A)**, and similar trends were observed when multiple profiles were removed. These results indicate that each PRE profile provides non-redundant local structural information. Consistently, conformational landscapes derived from simulations with fewer PRE profiles (**Figure 4D**) progressively lose the double-well feature observed with the full dataset, highlighting the necessity of multiple PRE profiles for resolving conformational heterogeneity.

Finally, we examined how many replicas are required to capture structural heterogeneity and reproduce ensemble-averaged experimental observables in parallel simulations. Although increasing the number of replicas can improve agreement with experiment, such gains are expected to plateau, while incurring increasing computational cost. To identify a practical balance, we performed simulations with 1, 4, 8, 16, and 32 replicas under identical restraint parameters. As shown in **Figure 4A**, improvements beyond 16 replicas were minimal, indicating diminishing returns with further increases. The corresponding conformational landscapes (**Figure 4E**) support this conclusion: simulations with few replicas are confined to one or a small number of restraint-averaged conformations, suppressing heterogeneity, whereas a sufficient number of replicas allows the system to explore the full conformational space under experimental restraints.

Taken together, these results clarify why resolving structural heterogeneity in IDPs requires both orthogonal experimental restraints and explicit ensemble sampling. Posterior ensemble reweighting is fundamentally limited by the conformational diversity of the initial simulation pool and cannot recover conformations that are not already sampled, leading to poor agreement with fine PRE data. SAXS and PRE provide complementary global and local information, and neither dataset alone is sufficient to define the full conformational landscape. Moreover, multi-replica simulations are essential for preserving heterogeneity under experimental restraints, as insufficient replica numbers drive the system toward artificial restraint-averaged conformations. By integrating orthogonal experimental measurements within a multi-replica framework, MARS overcomes these limitations and enables consistent reconstruction of both dominant and low-populated conformational states of ERα-NTD. A detailed summary of computational benchmarks is provided in **Table S2**.

## Conclusion

In this work, we introduced MARS, a Multi-replica Averaged Restraint Simulation framework for constructing experimentally consistent conformational ensembles of IDPs. By explicitly accounting for the ensemble-averaged nature of experimental observables, MARS enables simulations to reconcile heterogeneous conformational states with orthogonal experimental data. Applied to ERα-NTD, MARS integrates SAXS measurements and six PRE profiles to generate a structural ensemble that quantitatively reproduces the experimental observations. These results demonstrate that, heterogeneous IDP conformational ensembles can be constructed with high accuracy when a well-designed computational strategy is integrated with orthogonal experimental data.

The conformational ensemble of ERα-NTD revealed a landscape comprising two distinct conformational states. Notably, the low-populated compact state, stabilized by inter-cluster interactions that are difficult to infer from ensemble-averaged experiments alone, is consistent with previously determined phosphorylation-dependent activation mechanisms.^53^ These results illustrate the ability of MARS to uncover functionally relevant, low-populated conformational states that are obscured by ensemble-averaged observables.

In addition, we systematically dissected the methodological requirements for resolving IDP conformational heterogeneity. Although traditional ensemble reweighting or selection methods have achieved notable success in many systems^38–45^, its applicability can be limited in cases such as ERα-NTD, where low-populated compact states are difficult to sample. Examination of simulations restrained independently by SAXS or PRE data highlights the strong complementarity of global and local experimental information and demonstrates that neither dataset alone is sufficient to define the conformational landscape. Jackknife analysis of PRE restraints further reveals that even hundreds of distance-sensitive measurements provide largely non-redundant information for the conformational breadth of a flexible IDP. Finally, replica-number analysis establishes that multi-replica sampling is essential for preserving structural heterogeneity under experimental restraints, with convergence achieved once the ensemble has sufficient freedom to represent multiple coexisting conformational states.

Although MARS incurs higher computational cost than reweighting, its ability to generate missing conformations and consistently integrate orthogonal experimental datasets makes it well suited for modern IDP characterization. As the diversity and resolution of experimental measurements for disordered proteins continue to expand, MARS provides a general and scalable strategy for constructing mechanistically informative ensembles. More broadly, this work highlights an open challenge for the field: determining how experimental information content, sampling strategy, and model resolution jointly govern ensemble convergence and the resolvability of fine-grained interactions in intrinsically disordered systems.

## Methods

### Multi-replica Averaged Restraint Simulations

Within the MARS framework, we generated experimentally consistent ensembles by sequentially incorporating PRE and SAXS restraints into multi-replica simulations via observable-specific biasing potentials. The functional forms of the biasing potentials are as follows:

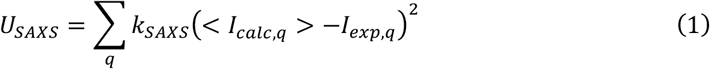

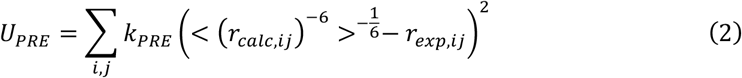

where <…> stands for the average over conformational ensembles. Here, *I*_*q*_ is the SAXS intensity at q; *i* represents the PRE labeling positions, and *j* denotes the residue for which the PRE intensity ratio relative to the label is measured; *r*_*exp,ij*_ is the distance back calculated from the PRE experiment data. The average over (*r*_*calc,ij*_)^−6^ is equivalent to average the relaxation enhancement 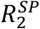.^64,65,42,66,67^ Distance between label *i* and residue *j* is calculated using the distance between the side chain bead of label *i* and the backbone bead of residue *j*.

Molecular dynamics simulations were performed using GROMACS 2023^68^ with a leap-frog integration algorithm and a time step of 20 fs. The starting configurations were randomly selected from an ERα-NTD ensemble in a previous work^69^. The initial equilibration was performed for 200 ps at 300 K using a Berendsen thermostat (couple time of 2 ps) and barostat (coupling time of 12 ps).^70^ Production simulations were then carried out for 500 ns at 300 K using a velocity-rescaling thermostat^71^ (coupling time of 1 ps) and at 1 bar using a Parrinello–Rahman barostat^72^ (coupling time of 12 ps). Van der Waals interactions were truncated at 11 Å, and Coulomb interactions were treated using the reaction-field method^73^ with the same cutoff. For productive simulations, the first 100 ns were discarded as equilibration and the remaining 400 ns were used for subsequent analysis.

For SAXS, the deviations between simulation and experiment are quantified using *χ*^2^ defined as:

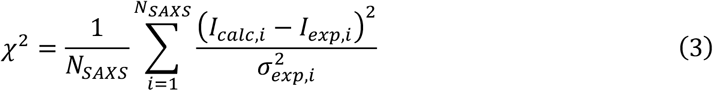

where *N*_*SAXS*_ is the number of SAXS q-values measured, *I*_*calc*_ is the intensity calculated from the simulated ensemble, *I*_*exp*_ is the experimental intensity, and σ_*exp*_ is the experimental uncertainty. For PRE, the deviations are quantified using the *Q*-factor defined as:

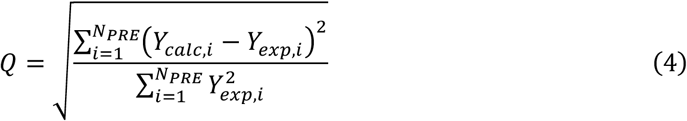

where *N*_*PRE*_ is the number of PRE intensity ratios measured, *Y*_*calc*_ *is t*he intensity ratio calculated from the simulated ensemble, and *Y*_*exp*_ *is th*e experimental intensity ratio.

Because MARTINI2 was originally parameterized for folded proteins, we first adjusted the protein–water interaction strength to reproduce the experimental *R*_*g*_ in unrestrained simulations (**Figure S1**), following recent recommendations^74^. We then optimized the SAXS and PRE biasing strengths. We incorporated the PRE restraints and systematically tuned their strength, selecting the value at the turning point beyond which further increases no longer improved agreement with the experimental data (**Figure S2A**). The SAXS restraints were then introduced while keeping the PRE restraint strength fixed at this optimized value. The same strategy was used to determine the optimal SAXS restraint strength (**Figure S2B**).

### Calculation of SAXS and PRE measurements from simulations

The calculation of the SAXS intensity is performed with the SAXS module in PLUMED 2.10^75^. For computational efficiency, a subset of points from the SAXS intensity profile was used as restraints, sampled with a stride of 10; the selected data points are listed in **Table S3**. When quantifying the deviations from the experimental data using the *Q* factor, all SAXS data points are included.

The calculation of PRE ratio is realized by first calculating the distance between the side chain bead of the labeled residue and the backbone bead of the residue of interest. Then the PRE ratio is calculated as

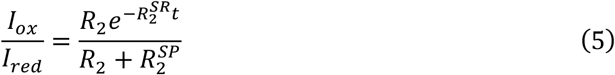

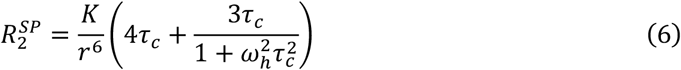

where *R*_2_ is the intrinsic relaxation of the backbone proton, 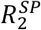 is the contribution to the relaxation caused by the paramagnetic label, *K* is a constant (1.23 × 10^-32^ cm^6^ sec^-2^) that describes the spin properties of the MTSL spin label, ω_*h*_ is the Larmor frequency of the proton spin (8.5 × 10^8^ sec^-1^), τ_*c*_ is the apparent PRE correlation time (3.27 × 10^-10^ sec), and *r* is the distance calculated from the simulation.^76^ As mentioned in the previous section, the average SAXS intensity is calculated directly over the intensities, while the average PRE ratio is obtained by first taking the average over 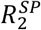 or equivalently *r*^−6^ and then calculating the corresponding PRE ratio.

### Time-lagged Independent Component Analysis (TICA)

TICA is a dimensionality-reduction method that identifies slow dynamical processes in molecular simulations by maximizing time-lagged autocorrelations of input features.^58^ Unlike PCA, which maximizes variance, TICA explicitly incorporates temporal evolving information. In practice, the simulation trajectories were first transformed from Cartesian coordinates into internal invariant features, such as bond lengths, bond angles, and pairwise contacts or distances. The covariance matrix was then computed using the mean-free features

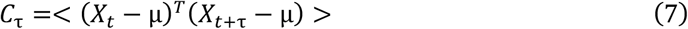

where <…> denotes the time average. TICA solves the generalized eigenvalue problem

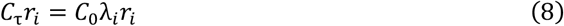

where the eigenvectors *r*_*i*_ define linear combinations of input features known as TICA coordinates, and the eigenvalues *λ*_*i*_ correspond to their normalized autocorrelations at lag time τ. The resulting TICA coordinates maximize slow motions and thus provide a low-dimensional representation of the essential conformational changes. In this study, we used pairwise distances between residues at odd-numbered positions, sampled at every other residue along the sequence, as input features. The analysis was performed using the TICA implementation embedded in the Deeptime Python module.^77^

### Ensemble reweighting

For the ensemble reweighting method tested in this study, we first performed a 16-replica simulation using the same MARTINI force field (see previous Method section and **Figure S1**) without experimental bias to generate the initial conformational pool. All structures in the pool were initially assigned equal weights. The objective function used for reweighting was defined as

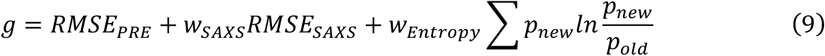

where *w*_*SAXS*_ and *w*_*Entropy*_ are the weighting coefficients for the SAXS RMSE and the entropy term, respectively; *p*_*new*_ represents the updated weights of the structures after perturbation, and *p*_*old*_ the weights before perturbation. During reweighting sampling at each step, the weights were randomly perturbed within the range of ±10% at each step. A change was accepted only if it reduces the value of the objective function. The parameters *w*_*SAXS*_ and *w*_*Entropy*_ were systematically varied to identify the optimal combination that minimized errors for both SAXS and PRE datasets while maintain a minimal deviation from the original pool to avoid overfitting (**Figure S7**).

## Acknowledgments

The authors acknowledge the Researching Computing at Arizona State University.

## Funding

The work was supported by the NIH R35GM146814 (WZ) and R01GM114056 (SY).

## Data and materials availability

The scripts are shared at https://github.com/wzhenglab/mars.

## Notes

### Competing Interest Statement

The authors have declared no competing interest.

