## Supplementary figures and tables. for "Resolving Conformational Heterogeneity in Intrinsically Disordered Proteins via Experimentally Guided Multi-Replica Simulations"

### Supporting Figures

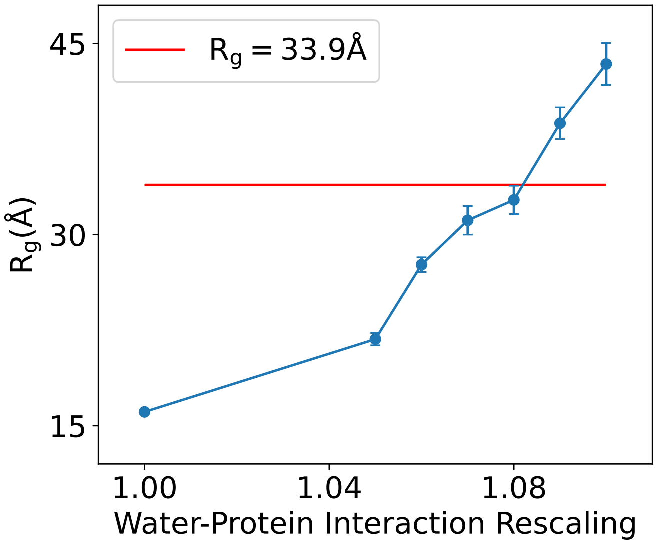


Figure S1: **ERα–NTD $R_g$ at Varying Protein–Water Interaction Strengths.** Blue dots represent the mean $R_g$ values of ERα–NTD computed at different protein–water interaction strengths. The red horizontal line denotes the $R_g$ inferred from SAXS data.

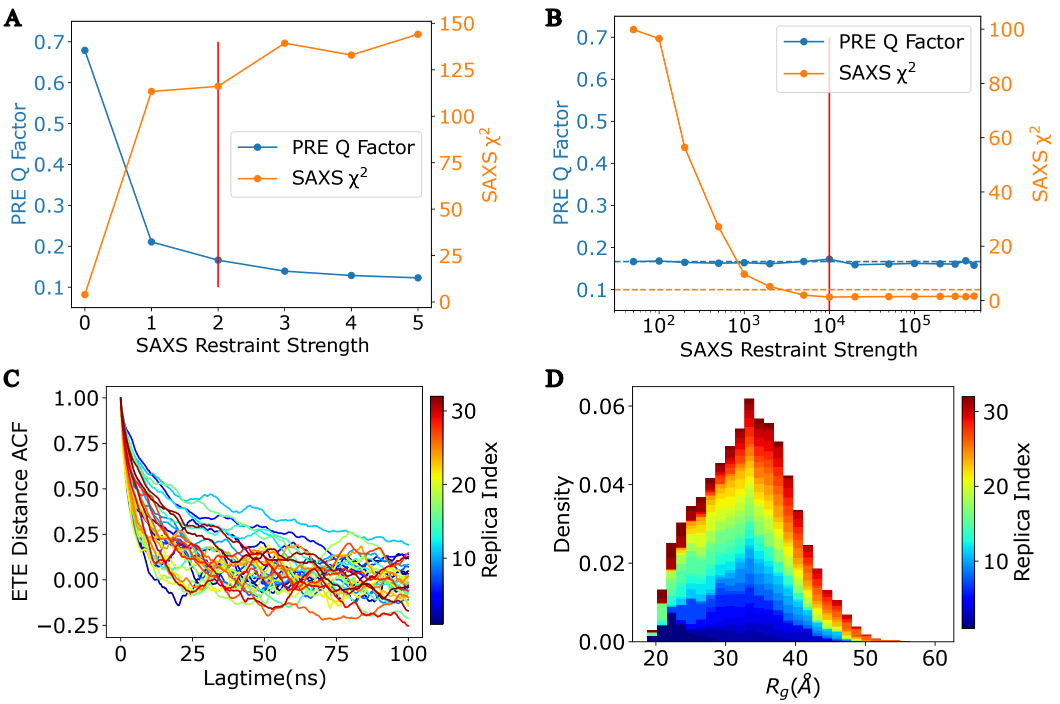


Figure S2: **Restraint Strength Tuning and Simulation Convergence Assessment.** (A) SAXS $\chi^2$ and PRE Q-factor as a function of the PRE restraint strength in the absence of SAXS restraints. The red vertical line indicates the PRE restraint strength selected for subsequent simulations.
(B) SAXS $\chi^2$ and PRE Q-factor as a function of the SAXS restraint strength, with the PRE restraint strength fixed at the value chosen in (A). The red vertical line indicates the SAXS restraint strength selected for subsequent simulations. The blue dashed line denotes the PRE Q-factor from the PRE-only–restrained simulations at the selected strength, while the orange dashed line denotes the SAXS $\chi^2$ from the unrestrained simulations.
(C) Auto-correlation function (ACF) of the ER$\alpha$–NTD end-to-end (ETE) distance obtained from the 32-replica simulation performed using the restraint strengths selected in (A) and (B).
(D) Histogram of $R_g$ from the same 32-replica simulations.

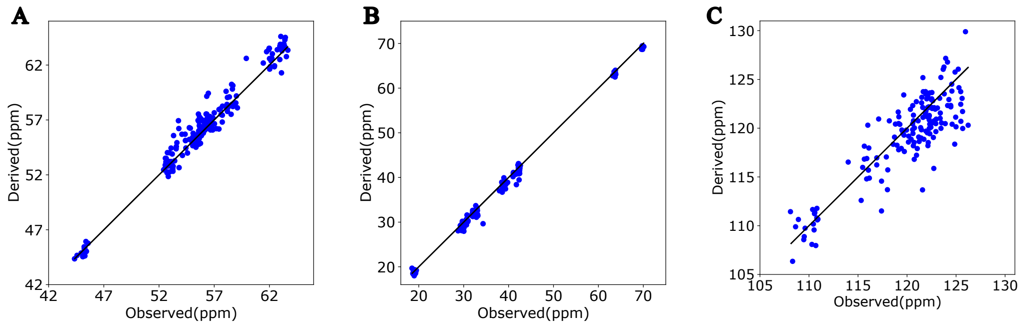


Figure S3: **Chemical Shift Comparison.**
Comparison of experimental and simulation-derived chemical shifts for (A) $^{13}C_\alpha$, (B) $^{13}C_\beta$, and (C) $^{15}N$.

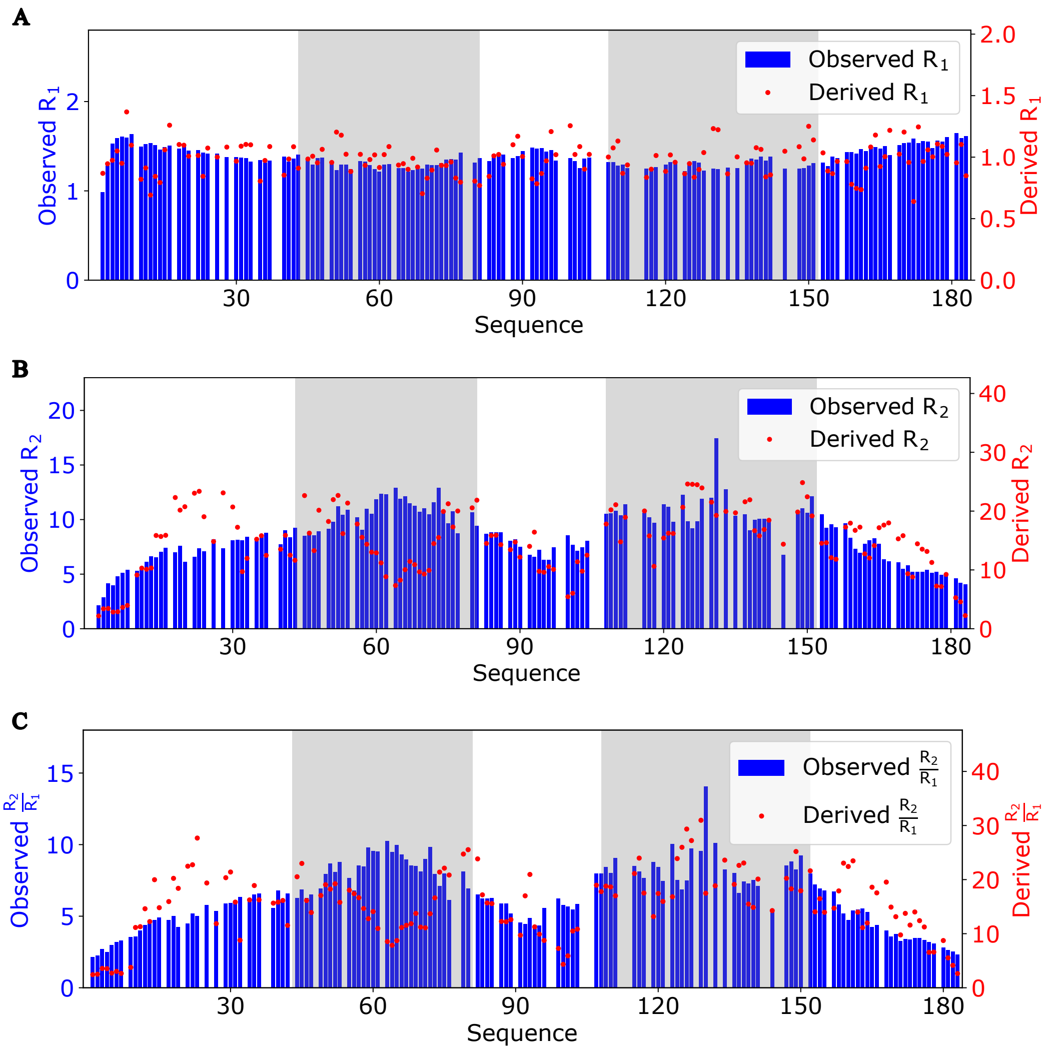


Figure S4: **NMR Relaxation Parameters.**
Experimental (blue) and simulation-derived (red) NMR relaxation parameters: (A) $R_1$, (B) $R_2$, and (C) $R_2/R_1$.

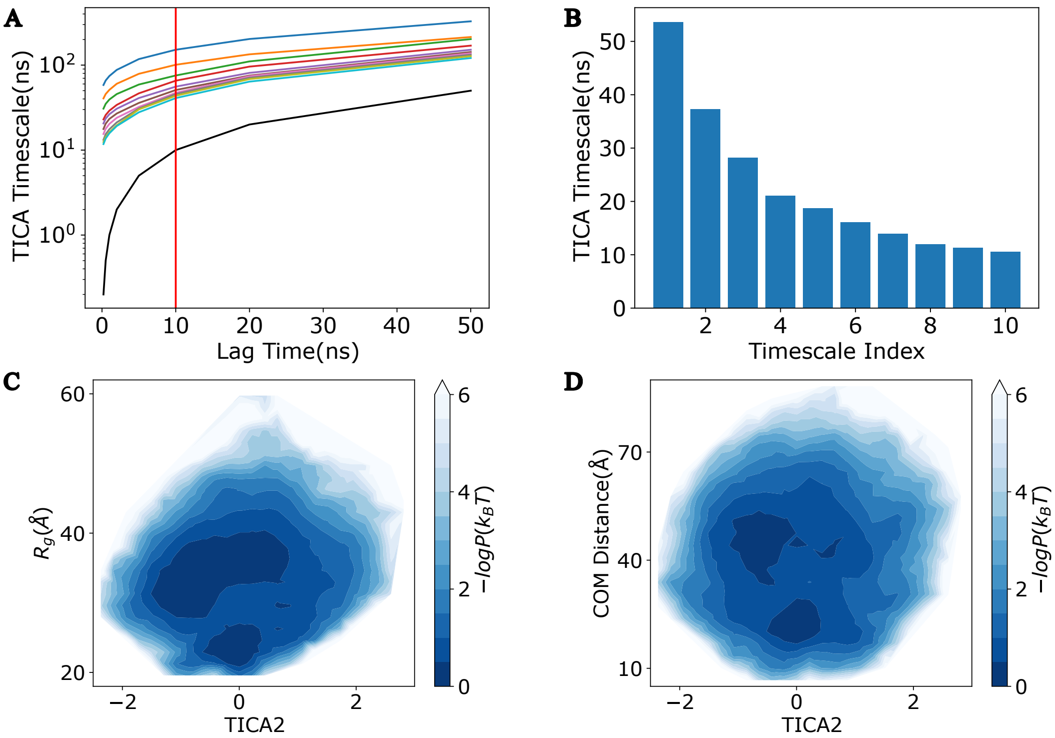


Figure S5: **TICA Timescales and Correlations between TICA2 and $R_g$, the COM Distance between two aromatic-rich sequence clusters (residues 43-81 and residues 108-152).**

(A) Convergence of TICA timescales as a function of the lag time. The red vertical line marks the lag time selected for subsequent analyses (10 ns).

(B) TICA timescale spectra evaluated at the lag time of 10 ns.

(C) Conformational landscape projected onto TICA2 and $R_g$.

(D) Conformational landscape projected onto TICA2 and the COM distance between the two clusters.

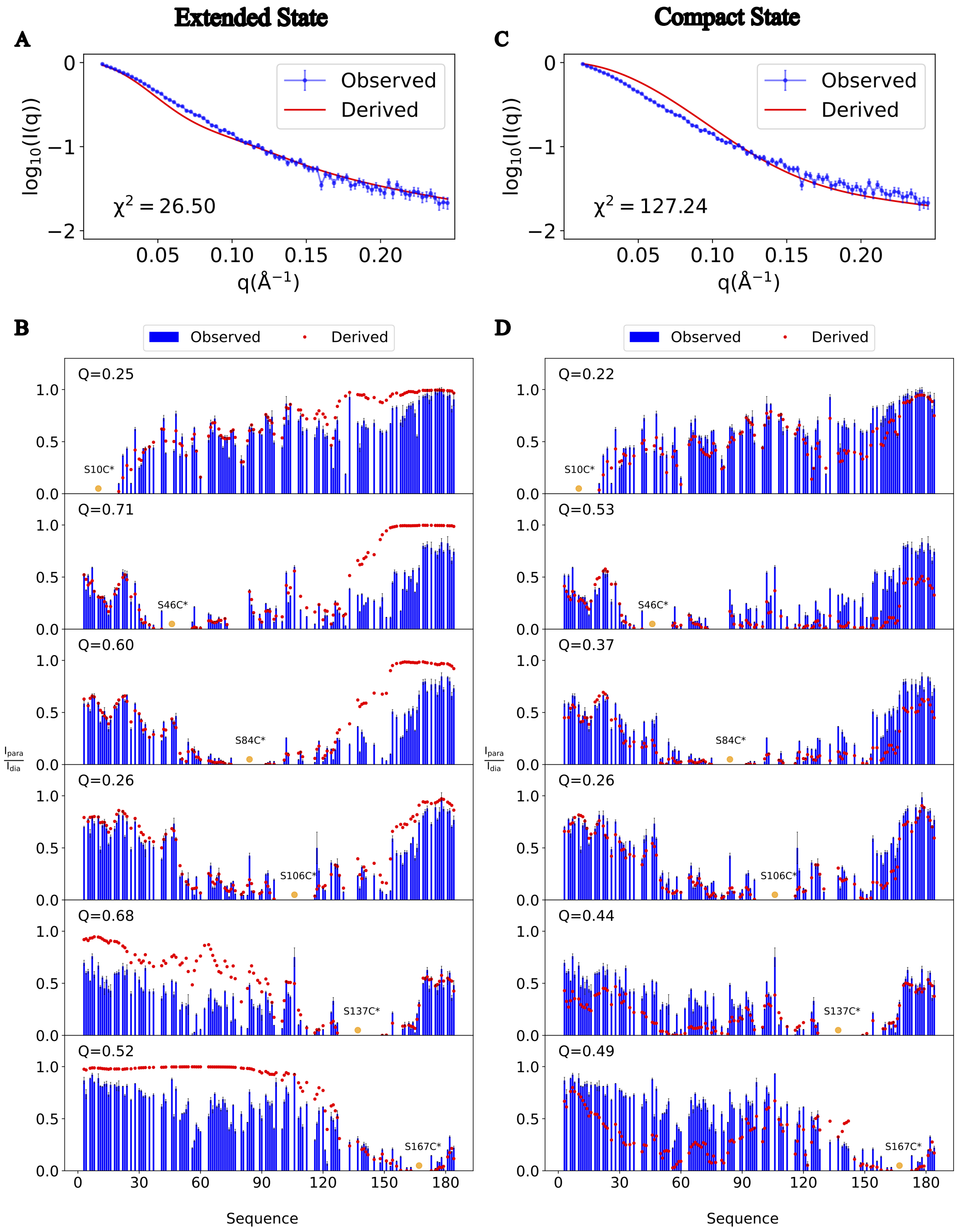


Figure S6: **Comparison of SAXS and PRE Experimental Data for the Two Conformational States.**
Experimental (blue) and simulation-derived (red) SAXS intensities for the extended state (A) and compact state (C), with corresponding $\chi^2$ values indicated in the figure legends. Experimental (blue) and simulation-derived (red) PRE profiles for the extended state (B) and compact state (D), with Q-factors reported in the figure legends. The spin-labeling position is highlighted in yellow.

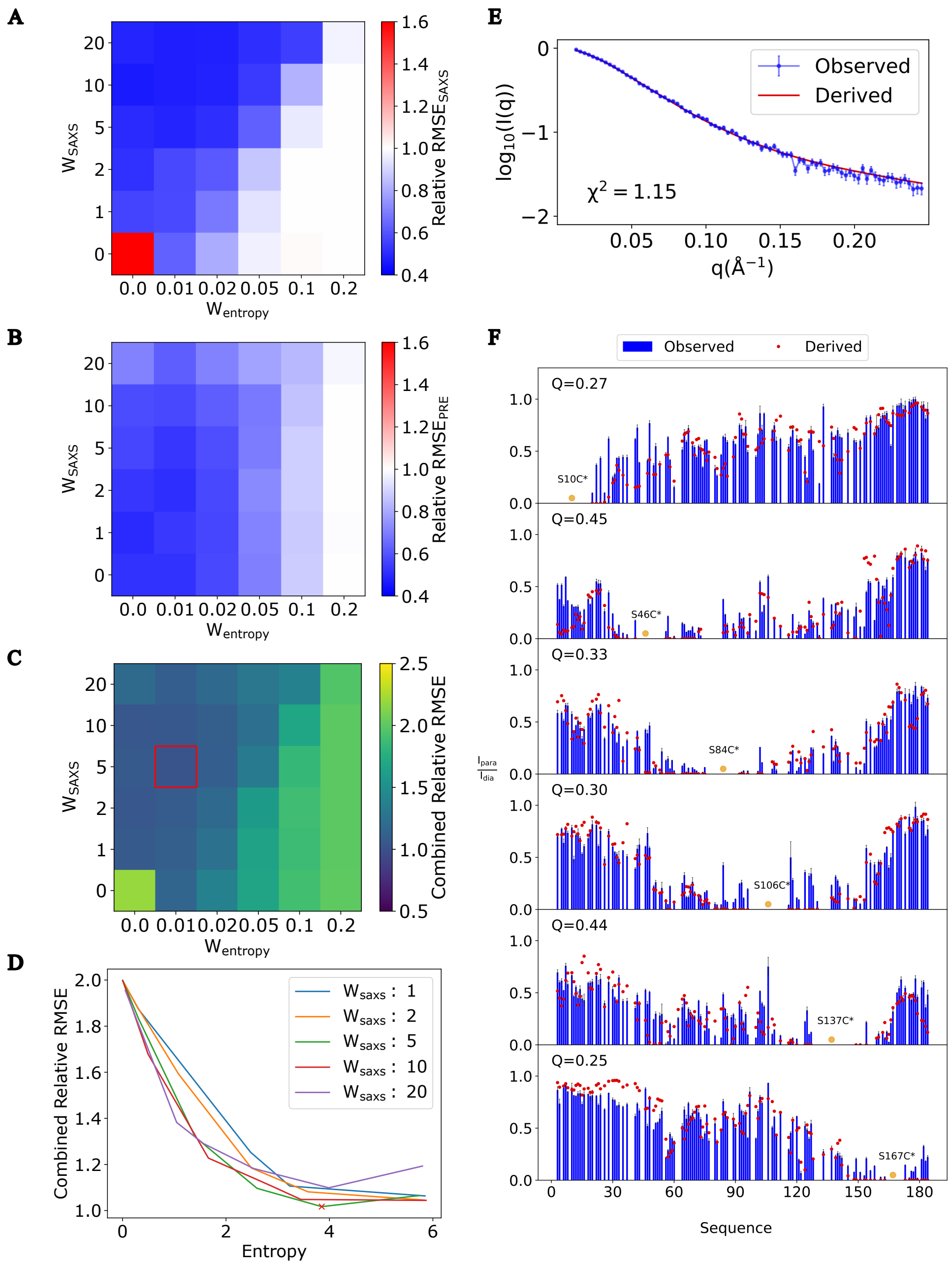


Figure S7: **Ensemble Reweighting Parameter Optimization and Comparison with SAXS and PRE Experimental Data.**

(A) Relative RMSE of SAXS evaluated at different weights assigned to the SAXS term and the entropy term in the objective function.

(B) Relative RMSE of PRE data under the same range of weighting schemes.

(C) Combined SAXS and PRE relative RMSE, with the red square indicating the optimal weight parameters selected for subsequent analyses.

(D) Combined relative RMSE plotted against the Shannon entropy for varying SAXS weights with the red cross indicating the result from the optimal weight parameters.

(E) Experimental (blue) and reweighted-ensemble–derived (red) SAXS intensities, with corresponding $\chi^2$ values noted in the figure legend.

(F) Experimental (blue) and reweighted-ensemble–derived (red) PRE measurements, with Q-factors reported in the figure legend.

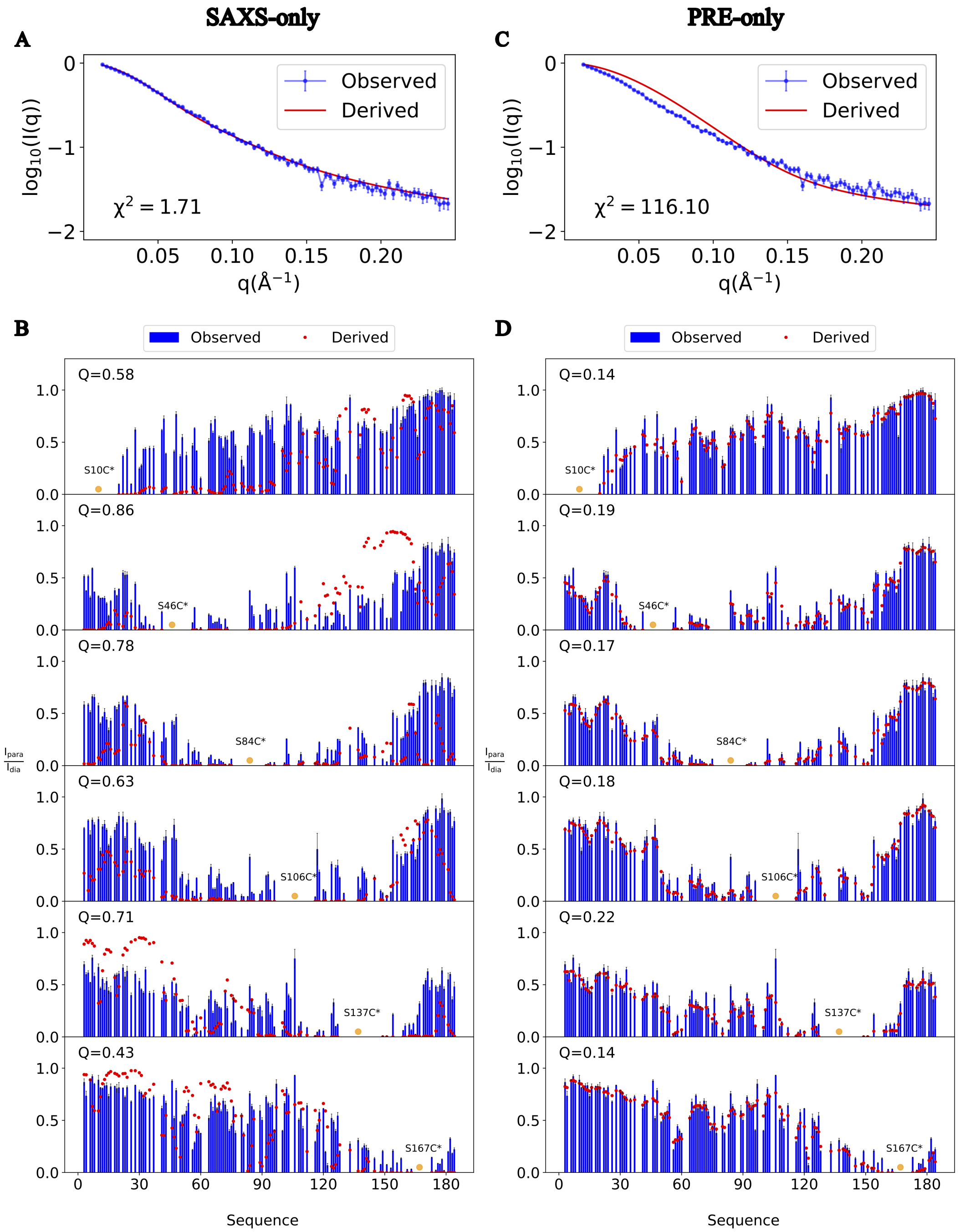


Figure S8: **Comparison of SAXS and PRE Experimental Data from Single-Source Restraint Simulations.**

Experimental (blue) and simulation-derived (red) SAXS intensities from the SAXS-restraint–only simulation (A) and the PRE-restraint–only simulation (C), with corresponding $\chi^2$ values indicated in the figure legends.

Experimental (blue) and simulation-derived (red) PRE profiles from the SAXS-restraint–only simulation (B) and the PRE-restraint–only simulation (D), with Q-factors noted in the figure legends. The spin-labeling position is highlighted in yellow.

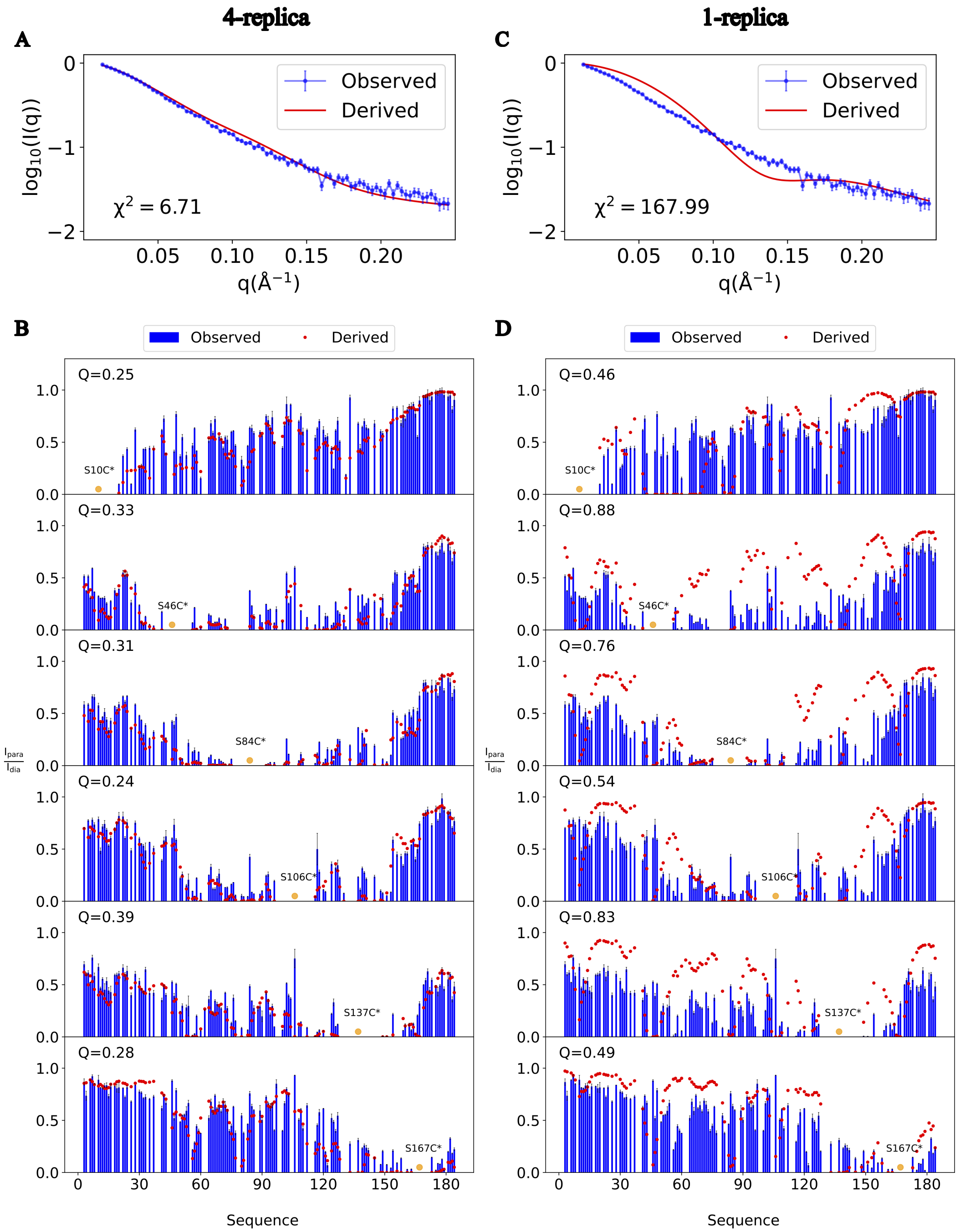


Figure S9: **Comparison of SAXS and PRE Experimental Data from Single-Source Restraint Simulations.**

Experimental (blue) and simulation-derived (red) SAXS intensities from the 4-replica simulation (A) and the single-replica simulation (C), with corresponding $\chi^2$ values indicated in the figure legends.

Experimental (blue) and simulation-derived (red) PRE profiles from the 4-replica simulation (B) and the single-replica simulation (D), with Q-factors noted in the figure legends. The spin-labeling position is highlighted in yellow.

### Supporting Tables

Table S1: Combinations Tested for Simulations with Part of PRE Restraints.

| Number of PRE Profile Included | Included PRE Profiles with Label Positions at |
|---|---|
| 1 | [10]; [46]; [84]; [106]; [137]; [167] |
| 2 | [10, 46]; [10, 84]; [46, 84]<br>[46, 106]; [46, 137]; [46, 167]<br>[84, 106]; [84, 137]; [84, 167] |
| 3 | [10, 46, 84]; [46, 84, 106]<br>[46, 84, 137]; [46, 84, 167] |
| 4 | [10, 46, 84, 106]; [10, 46, 84, 137]<br>[10, 46, 84, 167]; [10, 46, 106, 167]<br>[10, 46, 137, 167]; [10, 84, 106, 167]<br>[10, 106, 137, 167]; [46, 84, 106, 137]<br>[46, 84, 106, 167]; [46, 84, 137, 167] |
| 5 | [10, 46, 84, 106, 137]<br>[10, 46, 84, 106, 167]<br>[10, 46, 84, 137, 167]<br>[10, 46, 106, 137, 167]<br>[10, 84, 106, 137, 167]<br>[46, 84, 106, 137, 167] |

Table S2: Benchmarks of Computational Burden of Simulations under Different Restraint and Replica Settings.

| Simulation Settings | Simulation Length in 24h[ns] |
|---|---|
| Simulation with Both SAXS and PRE Restraints | |
| 32 Replicas | 95 |
| 16 Replicas | 96 |
| 8 Replicas | 99 |
| 4 Replicas | 102 |
| 1 Replica | 362 |
| Simulation with Only PRE Restraints | |
| 16 Replicas | 300 |
| 8 Replicas | 402 |
| 4 Replicas | 408 |
| 1 Replica | 471 |
| Simulation with Only SAXS Restraints | |
| 16 Replicas | 133 |
| 8 Replicas | 136 |
| 4 Replicas | 136 |
| 1 Replica | 367 |
| Simulation without Restraints | |
| 16 Replicas | 433 |
| 8 Replicas | 834 |
| 4 Replicas | 1704 |
| 1 Replica | 10435 |

Table S3: SAXS Data Points Used as Restraints.

| Scattering Vector [$\AA^{-1}$] | Experimental Intensity [a.u.] |
|---|---|
| 0.01262624 | 0.01171467 |
| 0.04099917 | 0.006806526 |
| 0.0693701 | 0.003258734 |
| 0.09773762 | 0.001785641 |
| 0.1261004 | 0.001053216 |
| 0.1544569 | 0.0006597623 |
| 0.182806 | 0.0004280769 |
| 0.211146 | 0.0004318829 |
